# Metagenomics untangles metabolic adaptations of Antarctic endolithic bacteria at the fringe of habitability

**DOI:** 10.1101/2023.07.30.551190

**Authors:** Claudia Coleine, Davide Albanese, Angelique E. Ray, Manuel Delgado-Baquerizo, Jason E. Stajich, Timothy J. Williams, Stefano Larsen, Susannah Tringe, Christa Pennacchio, Belinda C. Ferrari, Claudio Donati, Laura Selbmann

## Abstract

**Background:** Endolithic niches offer an ultimate refuge, supplying buffered conditions for microorganisms that dwell inside rock airspaces. Yet, survival and growth strategies of Antarctic endolithic microbes residing in Earths’ driest and coldest desert remains virtually unknown.

**Results:** From 109 endolithic microbiomes, 4,539 metagenome-assembled genomes were generated, 49.3% of which were novel candidate bacterial species. We present evidence that trace gas oxidation and atmospheric chemosynthesis may be the prevalent strategies supporting metabolic activity and persistence of these ecosystems at the fringe of life and the limits of habitability.

**Conclusions:** These results represent the foundation to untangle adaptability at the edge of sustainability on Earth and on other dry Earth-like planetary bodies such as Mars.

## Main

Permanently ice-free areas cover less than 1% of the Antarctic continent^1^ and include the coldest, driest and the most oligotrophic environments of Earth. Even so, Antarctic rocks are unexplored and isolated ecosystems that support highly diverse microbial communities; in such regions, highly adapted life forms subjected to a combination of poly-stresses still perpetuate^2,3^. Endolithic niches offer an ultimate refuge, supplying buffered conditions for microorganisms that dwell inside rock airspaces^4^.

Endolithic communities constitute simple food webs of varying complexity. Lichen-associated or free-living chlorophycean algae and *Cyanobacteria* function as primary producers, whilst fungi and more heterotrophic bacteria and support key ecosystem services such as nutrient cycling, rock weathering, and proto-soil formation^5,6^. Recent scientific studies considerably advanced our understanding of endolithic microbial biodiversity, environmental preferences, and extraordinary resistance to multiple stresses^5,7–9^. However, despite a number of studies being conducted at the community level, we still lack the most basic knowledge of how Antarctic endoliths survive the challenging conditions. A comprehensive genome catalog is the necessary first step to clarify the metabolic features and capabilities of these microorganisms and to elucidate how they survive such harsh conditions. Learning more about life under the extreme conditions is critical towards defining the fringe of habitability on Earth^10^.

To address this knowledge gap, we conducted a field survey including 109 endolithically colonized rocks, covering a wide plethora of regions and environments found in ice-free Antarctica, which includes a broad range of geo-environmental (e.g. altitudinal gradient, different rock typologies) and geographical distributions (i.e. Antarctic Peninsula, Northern Victoria Land, and McMurdo Dry Valleys; Figure 1a-c; Supplementary Table S1). We herein present the first Antarctic Rock Genomes Catalog (ARGC), which is the most comprehensive resource of bacterial metagenome-assembled genomes (MAGs) from terrestrial Antarctica to date.

**Figure 1.**
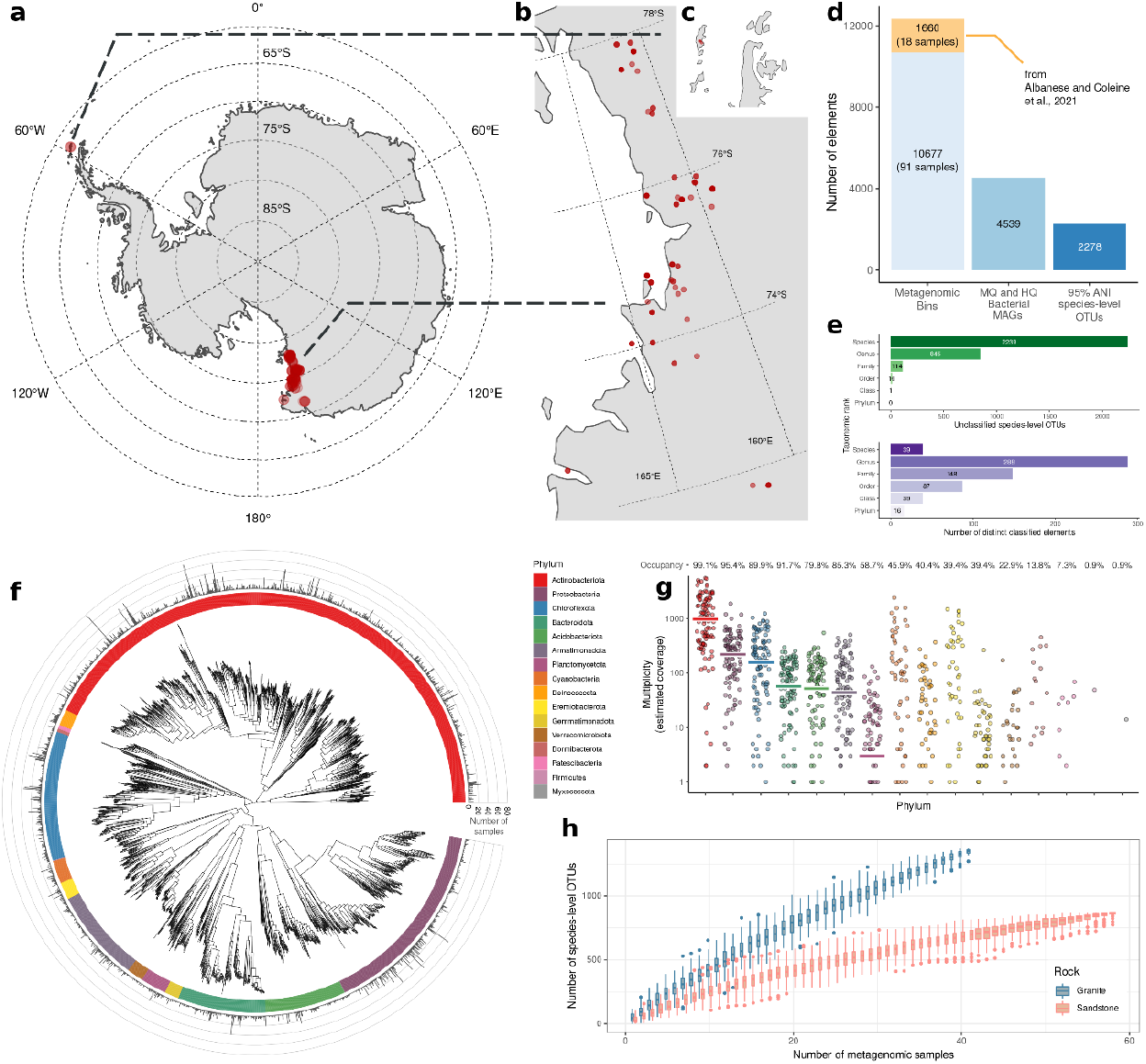
**a-c**, Map of Antarctica (a) and sampling sites (Victoria Land, b; Peninsula, c) (red dots). **d**, Number of MAGs and their quality-based classification. **e**, Upper bar plot: number of unclassified OTUs. Bottom bar plot: number of species, genera, families, orders, classes and phyla. **f**, Phylogenetic tree of the 2,278 OTUs built from the multiple sequence alignment of 120 GTDB marker genes. Barplot in the outer circle indicates the number of samples in which each OTUs was found. **g**, Phylum-level Mash Screen multiplicity for each sample, indicating sequence coverage. Horizontal lines represent the median values. The occupancy value indicates the percentage of samples that contains the underlying phylum. **h**, Number of OTUs as a function of the number of rock samples.

Following quality filtering (see Online Methods), 2,636 high-quality (HQ with ≥90% completeness and <5% contamination) and 1,903 medium-quality (MQ with ≥50% completeness and <10% contamination) bacterial MAGs were classified (Figure 1d; Supplementary Table S2, Supplementary Figures S1-5). The ARGC provides a complete picture of sandstone microbiomes across Antarctica, as revealed by the accumulation curves, which indicate that most species were retrieved; whilst, diversity in granite require further elucidation (Supplementary Figure S5). MAGs were then grouped at 95% average nucleotide identity (ANI) into 2,278 species-level bacterial operational taxonomic units (OTUs) (Figure 1e, f), 8.6 times more than previously reported^8^. All the OTUs can be assigned to known phyla, while 2,277, 2,262, 2,164 (95%), and 1,433 (63%) to known classes, orders, families and genera, respectively. Notably, 98.3% of species-level OTUs were distinct from the Genome Taxonomy Database (GTDB) reference genomes, representing 2,239 new candidate species (Figure 1e; Supplementary Table S3). On a phyla level, *Actinobacteriota* and *Proteobacteria* were dominant, with many new genomes of *Acidobacteriota, Chloroflexota*, and *Bacteroidota* also uncovered. *Actinomycetia* and *Thermoleophilia, Alphaproteobacteria*, and *Chloroflexia* classes were the most abundant and recurrent in the dataset (Figure 1g, Supplementary Figure S6; Supplementary Tables S4, S5). The dominant orders were *Mycobacteriales* (38%), *Actinomycetales* (15%), *Solirubrobacterales* (14%), *Acetobacterales* (12%), and *Thermomicrobiales* (7%) (Supplementary Table S6, S7).

To predict metabolic competencies, we retrieved 16,830,059 protein coding sequences (CDS) based on Prodigal analysis (see Methods). These CDS were dereplicated into 9,632,227, 6,997,885, 4,538,534 protein clusters using MMseqs2 with identity thresholds of 95%, 80% and 50% respectively. Moreover, 50% protein cluster representatives were searched against the UniProt Reference Clusters^11^ (UniRef, see Methods); since only 52.4% of the proteins displayed at least one match within the database, this resource should lay the foundation for future Antarctic terrestrial catalog.

During functional analysis, we focused on two widespread survival and growth strategies that allow microbiomes to persevere in extreme, oligotrophic environments; autotrophic metabolism, particularly trace gas chemosynthesis, and cold resistance adaptations. In cold edaphic deserts, energy generation through trace gas oxidation supports both microbial persistence and growth, with increased carbon fixation activity observed with aridity^12–14^. However, the significance of this strategy to endolithic microbiomes where photosynthetic microorganisms are more prevalent is questionable^15^.

High-affinity [NiFe]-hydrogenase genes, including forms 1h, 1l, 1m and 2a, are widely represented in our dataset, occurring in 41.1% of all dereplicated MAGs, including *Ca*. Dormibacterota (88.9%), *Eremiobacterota* (80.2%), *Actinobacteriota* (59.1%), *Gemmatimonadota* (57.1%), *Chloroflexota* (53.0%), *Acidobacteriota* (43.9%), *Verrucomicrobiota* (25.8%), *Planctomycetota* (13.4%), *Cyanobacteria* (7.5%), *Bacteroidota* (7.3%), *Proteobacteria* (6.1%), and *Armatimonadota* (4.8%) (Figure 2). The oxidation of trace levels of hydrogen gas plays a key role for persistence in dormant state and is a wide-spread ability in both Bacteria and Archaea in terrestrial and marine ecosystems^16,17^. The same strategy may be therefore crucial to support endolithic microbiomes whose active metabolism is, as average, limited to 1,000 h per year only^18^.

**Figure 2.**
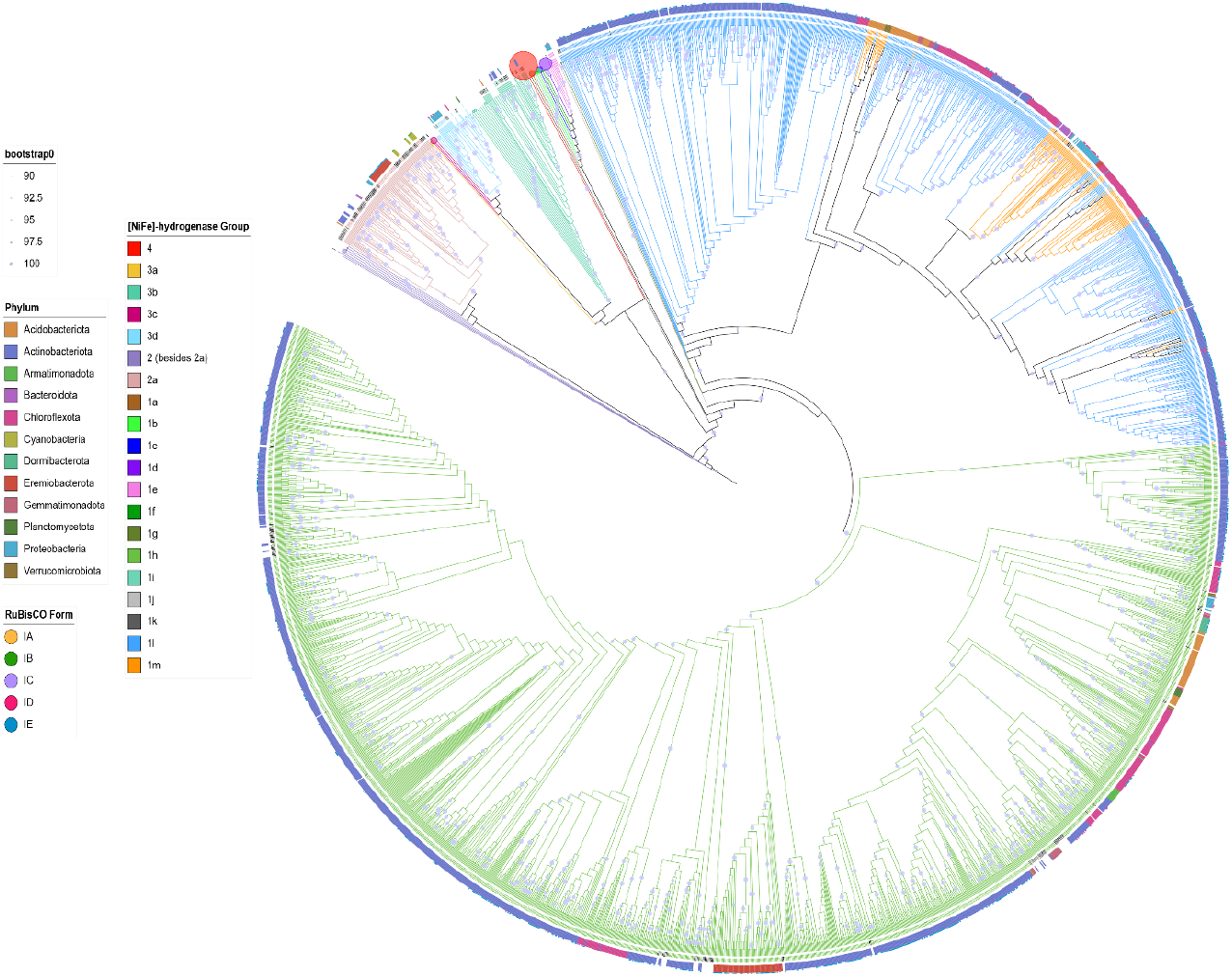
Maximum likelihood phylogenetic tree of [NiFe]-hydrogenase gene sequences obtained from our MAGs (n = 2433), with reference sequences obtained from the HydB and previous phylogenetic analysis. Branches and reference gene labels are colored according to the group of [NiFe]-hydrogenase. Bootstrap values >90% are depicted as filled circles on branches, with size reflecting value, and 1000 ultrafast bootstrap iterations applied. The phyla of the originating MAGs assembled in this study are displayed in a color-coded outer ring. In cases where RuBisCO large subunit gene/s co-occurred within these genomes, the proportion of forms present is indicated by external pie charts.

Autotrophic metabolisms are vital under such strict oligotrophic conditions and were indeed pervasive amongst the bacterial MAGs uncovered. Specifically, representatives from 7 of the 15 phyla presented signatures for carbon fixation. Phototrophic metabolism, mostly largely present in *Cyanobacteria*, is based on photolysis and requires water to take place. Evidence presented here suggests that trace gas oxidation may produce enough energy to not only support persistence but also to fuel the CBB cycle in a subset of the residing bacterial taxa, through the process of atmospheric chemosynthesis. This process is limited to cold soil deserts, while scarce to no carbon fixation activity has been observed yet in other environments^14,19^. Here we provide first-time evidence that atmospheric chemosynthesis could be extended to endolithic populations and may be a key adaptation for Carbon organization under highly dry conditions, with this process also proposed to be water-producing^20^. High-affinity [NiFe]-hydrogenases co-occurred alongside light-independent RuBisCO (1E/D) in 72.2% of *Ca*. Dormibacterota, 62.3% of *Eremiobacterota*, 20.6% of *Actinobacteriota*, 8.8% of *Chloroflexota*, 2.9% of *Gemmatimonadota* and 2.5% of *Proteobacteria* MAGs (Supplementary Figure S7), with RuBisCO form IE dominant accounting for 92.7% of those detected. These genetic indicators suggest that atmospheric chemosynthesis, as a fundamental process for primary production in hyper-arid cold environments, may be extended beyond soils to endolithic niches. RuBisCO form ID, showing a CO2 high affinity, is better adapted to a higher O2/CO2 ratio and requires less energetic or nutrient investment to attain high carboxylation rates; this finding suggests that, although uncommon, other RuBisCO forms may play a role in this chemoautotrophic process^21^. We propose that the plethora of RuBisCO forms found, displaying various efficiency, specificities, and affinities, enables the community to modulate its activity shifting from dormant to active state; this is paramount to adapt and exploit extreme and fluctuating microenvironments.

Aerobic respiration was predominant among endolithic MAGs (Supplementary Table S8; Figure 3); yet, the ability to use alternative e-acceptors via formate dehydrogenase, were limited to rare phyla, particularly in *Thermoanaerobaculia*, which was composed of one single family of anaerobic bacteria. The presence of additional chemosynthetic pathways, alternative to atmospheric chemosynthesis, using e-donors via Arsenate reductase were also found in a few (7) phyla, particularly abundant in *Bacilli*. This plethora of abilities to exploit various e-donors or acceptors increase the possibility of adaptability and survival of the whole community.

**Figure 3.**
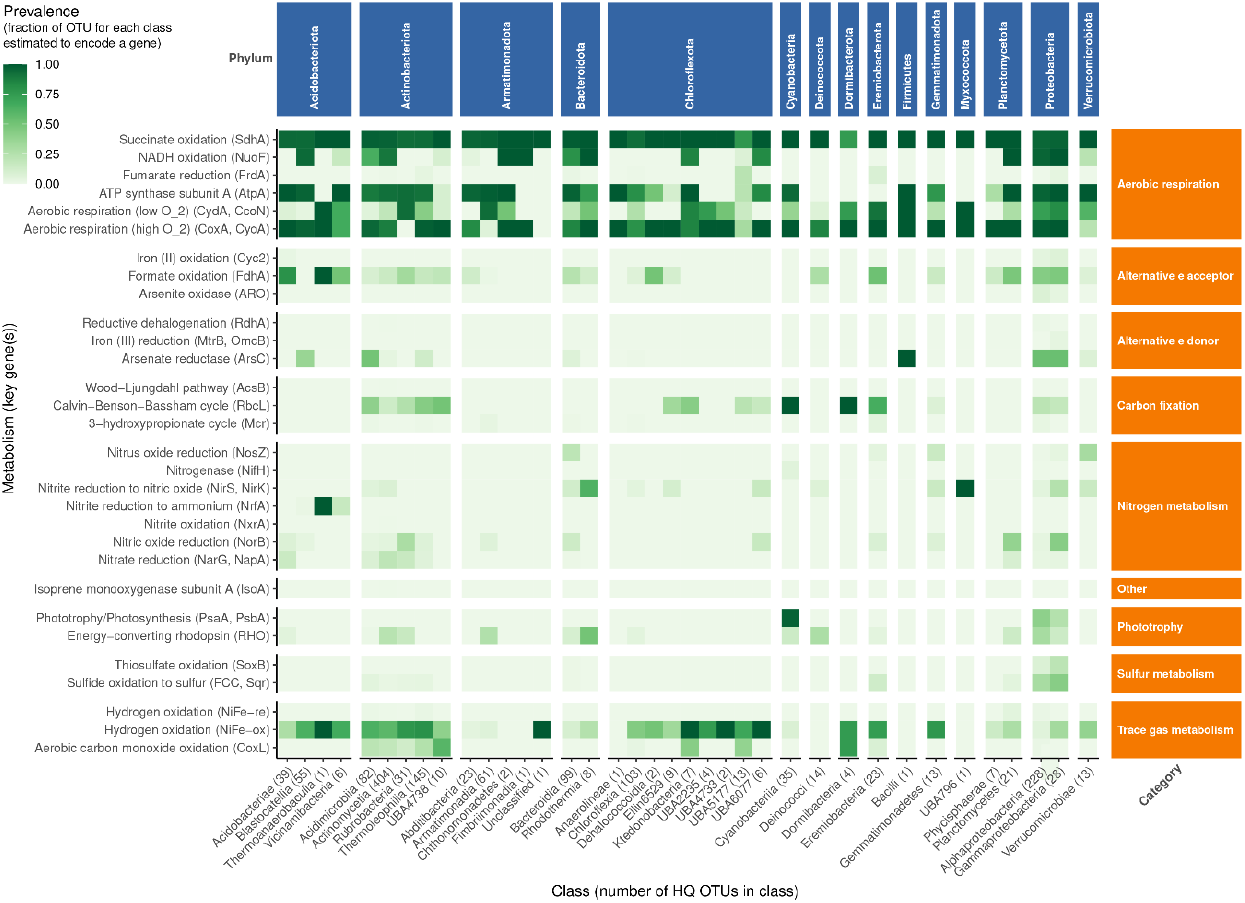
Metabolic potential of the species-level OTUs in Antarctic endolithic communities. The squared green cells represent the proportion of HQ OTUs in each class estimated to encode a particular metabolism. The analysis includes 1503 HQ OTUs partitioned in 37 classes and 15 phyla (blue rectangles), encompassing 30 key metabolisms partitioned in 9 categories (orange rectangles). NiFe-re and NiFe-ox indicates NiFe hydrogenases involved in H_2_ production (groups 3 and 4) H_2_ oxidation (groups 1 and 2a) respectively.

Lastly, below-freezing temperatures are a main challenge to life that can influence metabolic activity; reaching temperatures as low as −89°C, Antarctica is the coldest continent on the planet. We found that Antarctic endolithic bacteria encompass an innate adaptive capacity to cope with life in the persistent cold and the associated stresses. In fact, well-established genes involved in cold adaptation such as anti-freezing proteins (AFPs; e.g. 05934, K03522, K02959, K02386, K01993, K01934, K00658, K00627, K00324) were ubiquitous in all rock typologies and across all sampled areas (Supplementary Figure S8). This highlights the pivotal role of cold adaptation for survival at temperatures below 0°C^22,23^.

## Conclusions

Our study provides novel insights on the diversity of endolithic bacterial taxa thriving in the prohibitive conditions of Antarctica, and further identified survival strategies supporting their endurance at the limit of habitability. This resource represents the largest effort to date to capture the breadth of bacterial genomic diversity from Antarctic rocks. For the first time, we also unearthed the key and targeted adaptation strategies that allow microbes to spread and perpetuate in the harshest ecosystems. These results represent the foundation to untangle adaptability at the edge of sustainability on Earth and on other dry Earth-like planetary bodies such as Mars. This is also critical to inform us on the fate of microbial life in a warming and drying world.

## Declarations

### Ethics approval and consent to participate

Not applicable.

### Consent for publication

Not applicable.

### Availability of data and materials

Metagenomes raw data are available under the NCBI accession numbers listed in Supplementary Table 9. MAGs and annotations for high-quality MAGs are available at the zenodo repository (DOI: 10.5281/zenodo.7313591).

## Acknowledgements

C.C. is supported by the European Commission under the Marie Sklodowska-Curie Grant Agreement No. 702057 (DRYLIFE). C.C. and L.S. wish to thank the Italian National Program for Antarctic Research for funding sampling campaigns and research activities in Italy in the frame of PNRA projects. The Italian Antarctic National Museum (MNA) is kindly acknowledged for financial support to the Mycological Section of the MNA and for providing rock samples used in this study stored in the Culture Collection of Antarctic fungi (MNA-CCFEE), University of Tuscia, Italy. M.D-B. is supported by a project from the Spanish Ministry of Science and Innovation (PID2020-115813RA-I00), and a project of the Fondo Europeo de Desarrollo Regional (FEDER) and the Consejería de Transformación Económica, Industria, Conocimiento y Universidades of the Junta de Andalucía (FEDER Andalucía 2014-2020 Objetivo temático ‘01 – Refuerzo de la investigación, el desarrollo tecnológico y la innovación’) associated with the research project P20_00879 (ANDABIOMA). J.E.S. is a CIFAR fellow in the Fungal Kingdom: Threats and Opportunities program. B.C.F. acknowledges support from the Australian Research Council Discovery Project (DP220103430). Part of this work (proposal 10.46936/10.25585/60000791) was conducted by the U.S. Department of Energy Joint Genome Institute (https://ror.org/04xm1d337), a DOE Office of Science User Facility, supported by the Office of Science of the US Department of Energy under Contract No. DE-AC02-05CH11231.

## Contributions

C.C., D.A., A.E.R., B.C.F., C.D., and L.S., designed the study; C.C. performed DNA extraction and quality check control; D.A., C.C., and A.R. analyzed the data; C.C., D.A., A.R., B.C.F., C.D., and L.S., interpreted the results and wrote the paper with input from all authors. The authors read and approved the final manuscript.

