## Supplementary material for "Metagenomics untangles metabolic adaptations of Antarctic endolithic bacteria at the fringe of habitability": Online Methods

### Study area

Rocks colonized by endolithic communities were collected in thirty-eight sites in Antarctica including Antarctic Peninsula (*n* = 3), McMurdo Dry Valleys, Southern Victoria Land (*n* = 27), and Northern Victoria Land (*n* = 79) during more than 20 years of Italian Antarctic Expeditions. Different rock typologies (sandstone *n* = 59, granite *n* = 43, quartz *n* = 5 , and basalt/dolerite *n* = 2) were collected along a latitudinal transect ranging from -62.10008 -58.51664 to -77.874 160.739 at different environmental conditions namely sun exposure (northern sun exposed and southern shady rocks) and an altitudinal transect from sea level to 3,100 m above sea level (a.s.l.) to provide a comprehensive overview of Antarctic endolithic diversity (Figure 1, Supplementary Table 1). The presence of endolithic colonization was assessed by direct observation in situ. Rocks were excised using a geologic hammer and sterile chisel, and rock samples, preserved in sterile plastic bags, transported, and stored at −20 °C in the Culture Collection of Antarctic fungi of the Mycological Section of the Italian Antarctic National Museum (MNA-FCC), until downstream analysis.

### DNA extraction, library preparation, and sequencing

Metagenomic DNA was extracted from 1 g of crushed rocks using DNeasy PowerSoil Pro Kit (Qiagen, German), quality checked by electrophoresis using a 1.5% agarose gel and Nanodrop spectrophotometer (Thermofisher, USA) and quantified using the Qubit dsDNA HS Assay Kit (Life Technologies, USA) according to Coleine et al. (2021). Shotgun metagenomic sequencing paired-end libraries were constructed and sequenced as 2×150 bp using the Illumina NovaSeq platform (Illumina Inc, San Diego, CA) at the Edmund Mach Foundation (San Michele all'Adige, Italy) and at the DOE Joint Genome Institute (JGI).

### Sequencing reads preparation, assembly and binning

The metashot/mag-illumina v2.0.0 Nextflow-based (Di Tommaso et al. 2017) workflow (https://github.com/metashot/mag-illumina, parameters: --metaspades_k 21,33,55,77,99) was used to perform raw reads quality trimming and filtering, assembly and contings binning on the 91 metagenomic samples. In brief, adapter trimming, contaminant (artifacts and and spike-ins) and quality filtering were performed using BBDuk (BBMap/BBTools v38.79, https://sourceforge.net/projects/bbmap/). During the quality filtering procedure i) raw reads were quality-trimmed to Q6 using the Phred algorithm; ii) reads that contained 4 or more "N" bases, had an average quality below 10, shorter than 50 bp or under 50% of the original length were removed. Samples were then assembled individually with SPAdes (Nurk et al. 2017) v3.15.1 (parameters --meta -k 21,33,55,77,99).

Metagenomic contigs were binned into candidate metagenome-assembled genomes (MAGs) using MetaBAT 2 (Metagenome Binning based on Abundance and Tetranucleotide frequency) (Kang et al. 2019) v2.12.1. Briefly, high-quality reads were mapped on assembled contigs using Bowtie2 (Langmead and Salzberg 2012) v2.3.4.3. Samtools (Li et al. 2009) (htslib v1.9) was used to create and sort the BAM files. The depth of coverage was estimated by applying the MetaBAT2 script "jgi_summarize_bam_contig_depths". Finally, contigs sequences and the depth of coverage estimates were used by MetaBAT2 to recover the 10,677 bins.

### Quality assessment, filtering and dereplication

The resulting bins were combined with the 1660 metagenomic bins from (Albanese et al. 2021) and analyzed using themetashot/prok-quality (Albanese and Donati 2021) v1.2.3 (parameters --gunc_filter --gunc_db gunc_db_2.0.4.dmnd) workflow. Briefly, completeness, redundant and non-redundant contamination (Orakov et al., n.d.) estimates were obtained by CheckM (Parks et al. 2015) v1.1.2 and GUNC (Orakov et al., n.d.). Bins with completeness estimates of <50%, more than 10% contamination and that did not pass the GUNC filter were discarded, resulting in a total of 4,540 filtered prokaryotic MAGs. MAGs were classified into "high-quality draft" (HQ) with >90% completeness and <5% contamination and "medium-quality draft" (MQ) with completeness estimates of ≥50% and less than 10% contamination. Species-level operational taxonomic units (OTUs) were identified by clustering HQ and MQ MAGs at 95% average nucleotide identity (ANI) using dRep (Olm et al. 2017) v2.6.2, resulting in a total of 2,279 OTUs. For each species-level OTUs, the MAG with the highest quality score was chosen as representative. The score was computed using the formula: score = completeness - 5 x contamination + 0.5 x log(N50) (Albanese and Donati 2021).

### Taxonomic classification of prokaryotic MAGs

Species-level OTUs representative MAGs were taxonomically classified using the metashot/prok-classify v1.2.1 workflow (https://github.com/metashot/prok-classify, parameters: --gtdbtk_db release202). The workflow includes the genome taxonomy database toolkit (GTDB-Tk) (Chaumeil et al. 2019) v1.5.0 and the GTDB release 202, following the recently proposed nomenclature of prokaryotes (Parks et al. 2022). A single OTU was classified as archaea and was removed from subsequent analyses. Approximately-maximum-likelihood phylogenetic tree from the GTDB protein alignments of the 2,278 bacterial OTU representatives was inferred using FastTree (Price, Dehal, and Arkin 2010) v2.1.11 (default parameters).

### Bacterial OTU coverage estimates in metagenomes

The metashot/containment v1.0.0 workflow (https://github.com/metashot/containment, parameters: --min_identity 0.95 --winner_takes_all --sketch_size 10000) was used to determine the presence of the reconstructed bacterial OTU in the 109 Antarctic samples. Briefly, for each metagenome we applied the Mash Screen algorithm (Ondov et al. 2019) (Mash v2.1) in order to calculate the containment score for each OTU (i.e., the estimate of the similarity of an OTU representative to a sequence contained within the metagenome), its p value and the OTU median-multiplicity, as a proxy for the OTU coverage. The Mash Screen algorithm demonstrated to be in good agreement with the mapping-and-consensus procedure described in (Albanese et al. 2021).

### Functional annotation of bacterial MAGs

Functional annotation was performed using the workflow metashot/prok-annotate (https://github.com/metashot/prok-annotate, commit da2d0bb, parameters: --run_eggnog --eggnog_db emapperdb-5.0.2). Input MQ and HQ bacterial MAGs (n=4,539) were processed as follows: (i) 16,830,059 translated coding DNA sequences (CDSs) were predicted using Prokka (Seemann 2014) v1.14.5 which in turn wraps the gene predictor Prodigal (Hyatt et al. 2010) and (ii) functionally annotated using EggNOG-mapper (Cantalapiedra et al. 2021) (v2.1.4, parameters -m diamond --itype protein) against the eggNOG Orthologous Groups (OGs) database (Huerta-Cepas et al. 2019) v5.0.2. The eggNOG database integrates functional annotations collected from several sources, including Gene Ontology (GO) terms, KEGG functional orthologs (Kanehisa et al. 2014) and COG categories (Tatusov et al. 2000). For each species-level OTU, a target gene/ortholog was marked as "present" if more or equal than 80% of the HQ genomes which belong to the OTU encoded that gene/ortholog. Translated CDSs were de-replicated at 95%, 80% and 50% identity and an alignment fraction threshold of 80% using MMseqs2 (Mirdita, Steinegger, and Söding 2019) v13-1 (ref. 52) with the parameters "easy-linclust -e 0.001 --min-seq-id [IDENTITY] -c 0.80". 50% protein cluster representatives were searched against the UniProt Reference Clusters (UniRef50, release 2022_01, 23-Feb-2022, http://www.uniprot.org) with an identity threshold of 50% using the MMseqs2's easy-search protocol (parameters -e 0.001 --min-seq-id 0.5 --cov-mode 2 -c 0.8). Moreover, translated CDSs (n=16,830,059) were searched against the "Greening lab metabolic marker gene databases" (Greening 2021) using an identity threshold of 50% (parameters: easy-search --min-seq-id 0.5 --cov-mode 2 -c 0.8). Best hits were further filtered for some marker gene according to (Chen et al. 2021): [NiFe]-hydrogenases, [FeFe]-hydrogenases, CoxL, AmoA, NxrA and NuoF were filtered at 60% identity threshold, AtpA, YgfK, HbsT, ARO, and PsbA at 70%, and PsaA at 80%. For each species-level OTU, a target gene was marked as "present" if more or equal than 80% of the HQ genomes which belong to the OTU encoded that gene.

### Phylogenetic Analysis of RuBisCO and [NiFe]-hydrogenase

A total 978 putative RuBisCO sequences and 2433 putative [NiFe]-hydrogenase sequences were yielded from the recovered MAGs. All sequences obtained were further classified into subforms using previously published databases and BLAST+ (ver. 2.12.0) (Camacho et al. 2009). In addition, separate phylogenetic analyses were conducted to visualize the forms of RuBisCO and [NiFe]-hydrogenase present. The extracted RuBisCO sequences were analyzed against 3129 reference sequences obtained through previous phylogenetic analysis of the Genome Taxonomy Database (Ray et al. 2022)(Ray et al., 2022). [NiFe]-hydrogenase sequences were analyzed against 2019 reference sequences obtained from the HydDB (Søndergaard, Pedersen, and Greening 2016)(Søndergaard et al., 2016) and previous phylogenetic analysis (Ray et al. 2022)(Ray et al., 2022).

Multiple sequence alignment was conducted using MAFFT (ver.7.407), applying the L-INS-i iterative refinement method (Katoh et al. 2002; Katoh and Standley 2013)(Katoh et al., 2002, Katoh and Standley, 2013). To remove poorly aligned regions, the resulting alignments were trimmed using trimAl (ver. 1.4.1), with a gap threshold of 0.5 (Capella-Gutiérrez, Silla-Martínez, and Gabaldón 2009)(Capella-Gutiérrez et al., 2009). Sequences with more than 50% gaps after alignment were removed. Maximum likelihood phylogenetic trees were constructed using IQ-Tree (ver. 1.6.10) (Nguyen et al. 2015)(Nguyen et al., 2015), applying 1000 ultrafast bootstrap iterations, hill-climbing nearest neighbor interchange (NNI) search and incorporating additional SH-like approximate likelihood ratio tests (SH-alrt) (Guindon et al. 2010)(Guindon et al., 2010). ModelFinder was used to determine the best evolutionary model (Nguyen et al. 2015), which was LG+F+R10 for the RuBisCO tree and LG+R10 for the [NiFe]-hydrogenase tree. Sequences that failed the chi2 test during tree building were removed.

The final consensus trees, comprising 1,347 RuBisCO sequences and 2,706 [NiFe]-hydrogenase sequences, were uploaded to Interactive Tree Of Life (iTOL) v6 (Letunic and Bork 2016)(Letunic and Bork, 2016) for visualization. Branches were color-coded according to the form of [NiFe]-hydrogenase or RuBisCO, and bootstrap values 90-100 were indicated by circles on the corresponding branches, with size corresponding to values. The phyla of the MAGs from which each sequence originated are displayed as a color-coded outer ring. In the [NiFe]-hydrogenase tree, if the RuBisCO large subunit co-occurred within the originating genome, then these sequences are marked by an outer pie chart, which depicts the proportion of RuBisCO forms detected. Complete trees showing all RuBisCO and [NiFe]-hydrogenase sequences are provided (Supplementary Information). Collapsed versions are also provided, focusing upon clades where sequences from the MAGs studied here were identified, namely RuBisCO form I and forms 1h, 1l, 1m, 1e, 2a, 3b, 3d [NiFe]-hydrogenase.

#### Downstream analysis

Downstream analysis was performed using the R environment (https://www.R-project.org/) v4.0.3 and the packages tidyverse v1.3.0, ggtree v2.4.1 and phytools v0.7-70.

5825–29. https://doi.org/10.1093/molbev/msab293.

Capella-Gutiérrez, Salvador, José M. Silla-Martínez, and Toni Gabaldón. 2009. “trimAl: A Tool for Automated Alignment Trimming in Large-Scale Phylogenetic Analyses.” *Bioinformatics*. https://doi.org/10.1093/bioinformatics/btp348.

Chaumeil, Pierre-Alain, Aaron J. Mussig, Philip Hugenholtz, and Donovan H. Parks. 2019. “GTDB-Tk: A Toolkit to Classify Genomes with the Genome Taxonomy Database.” *Bioinformatics* , November. https://doi.org/10.1093/bioinformatics/btz848.

Chen, Ya-Jou, Pok Man Leung, Jennifer L. Wood, Sean K. Bay, Philip Hugenholtz, Adam J. Kessler, Guy Shelley, et al. 2021. “Metabolic Flexibility Allows Bacterial Habitat Generalists to Become Dominant in a Frequently Disturbed Ecosystem.” *The ISME Journal*. https://doi.org/10.1038/s41396-021-00988-w.

Di Tommaso, Paolo, Maria Chatzou, Evan W. Floden, Pablo Prieto Barja, Emilio Palumbo, and Cedric Notredame. 2017. “Nextflow Enables Reproducible Computational Workflows.” *Nature Biotechnology* 35 (4): 316–19. https://doi.org/10.1038/nbt.3820.

Greening, Chris. 2021. “Greening Lab Metabolic Marker Gene Databases,” April. https://doi.org/10.26180/c.5230745.

Guindon, Stéphane, Jean-François Dufayard, Vincent Lefort, Maria Anisimova, Wim Hordijk, and Olivier Gascuel. 2010. “New Algorithms and Methods to Estimate Maximum-Likelihood Phylogenies: Assessing the Performance of PhyML 3.0.” *Systematic Biology* 59 (3): 307–21. https://doi.org/10.1093/sysbio/syq010.

Huerta-Cepas, Jaime, Damian Szklarczyk, Davide Heller, Ana Hernández-Plaza, Sofia K. Forslund, Helen Cook, Daniel R. Mende, et al. 2019. “eggNOG 5.0: A Hierarchical, Functionally and Phylogenetically Annotated Orthology Resource Based on 5090 Organisms and 2502 Viruses.” *Nucleic Acids Research*. https://doi.org/10.1093/nar/gky1085.

Hyatt, Doug, Gwo-Liang Chen, Philip F. Locascio, Miriam L. Land, Frank W. Larimer, and Loren J. Hauser. 2010. “Prodigal: Prokaryotic Gene Recognition and Translation Initiation Site Identification.” *BMC Bioinformatics* 11 (March): 119. https://doi.org/10.1186/1471-2105-11-119.

Kanehisa, Minoru, Susumu Goto, Yoko Sato, Masayuki Kawashima, Miho Furumichi, and Mao Tanabe. 2014. “Data, Information, Knowledge and Principle: Back to Metabolism in KEGG.” *Nucleic Acids Research* 42 (Database issue): D199–205. https://doi.org/10.1093/nar/gkt1076.

Kang, Dongwan D., Feng Li, Edward Kirton, Ashleigh Thomas, Rob Egan, Hong An, and Zhong Wang. 2019. “MetaBAT 2: An Adaptive Binning Algorithm for Robust and Efficient Genome Reconstruction from Metagenome Assemblies.” *PeerJ* 7 (July): e7359. https://doi.org/10.7717/peerj.7359.

Katoh, Kazutaka, Kazuharu Misawa, Kei-Ichi Kuma, and Takashi Miyata. 2002. “MAFFT: A Novel Method for Rapid Multiple Sequence Alignment Based on Fast Fourier Transform.” *Nucleic Acids Research* 30 (14): 3059–66. https://doi.org/10.1093/nar/gkf436.

Katoh, Kazutaka, and Daron M. Standley. 2013. “MAFFT Multiple Sequence Alignment Software Version 7: Improvements in Performance and Usability.” *Molecular Biology and Evolution* 30 (4): 772–80. https://doi.org/10.1093/molbev/mst010.

Langmead, Ben, and Steven L. Salzberg. 2012. "Fast Gapped-Read Alignment with Bowtie 2." *Nature Methods* 9 (4): 357–59. https://doi.org/10.1038/nmeth.1923.

Letunic, Ivica, and Peer Bork. 2016. "Interactive Tree of Life (iTOL) v3: An Online Tool for the Display and Annotation of Phylogenetic and Other Trees." *Nucleic Acids Research*. https://doi.org/10.1093/nar/gkw290.

Li, Heng, Bob Handsaker, Alec Wysoker, Tim Fennell, Jue Ruan, Nils Homer, Gabor Marth, Goncalo Abecasis, Richard Durbin, and 1000 Genome Project Data Processing Subgroup. 2009. "The Sequence Alignment/Map Format and SAMtools." *Bioinformatics* 25 (16): 2078–79. https://doi.org/10.1093/bioinformatics/btp352.

Mirdita, Milot, Martin Steinegger, and Johannes Söding. 2019. "MMseqs2 Desktop and Local Web Server App for Fast, Interactive Sequence Searches." *Bioinformatics* 35 (16): 2856–58. https://doi.org/10.1093/bioinformatics/bty1057.

Nguyen, Lam-Tung, Heiko A. Schmidt, Arndt von Haeseler, and Bui Quang Minh. 2015. "IQ-TREE: A Fast and Effective Stochastic Algorithm for Estimating Maximum-Likelihood Phylogenies." *Molecular Biology and Evolution*. https://doi.org/10.1093/molbev/msu300.

Nurk, Sergey, Dmitry Meleshko, Anton Korobeynikov, and Pavel A. Pevzner. 2017. "metaSPAdes: A New Versatile Metagenomic Assembler." *Genome Research* 27 (5): 824–34. https://doi.org/10.1101/gr.213959.116.

Olm, Matthew R., Christopher T. Brown, Brandon Brooks, and Jillian F. Banfield. 2017. "dRep: A Tool for Fast and Accurate Genomic Comparisons That Enables Improved Genome Recovery from Metagenomes through de-Replication." *The ISME Journal* 11 (12): 2864–68. https://doi.org/10.1038/ismej.2017.126.

Ondov, Brian D., Gabriel J. Starrett, Anna Sappington, Aleksandra Kostic, Sergey Koren, Christopher B. Buck, and Adam M. Phillippy. 2019. "Mash Screen: High-Throughput Sequence Containment Estimation for Genome Discovery." *Genome Biology* 20 (1): 232. https://doi.org/10.1186/s13059-019-1841-x.

Orakov, Askarbek, Anthony Fullam, Luis Pedro Coelho, Supriya Khedkar, Damian Szklarczyk, Daniel R. Mende, Thomas S. B. Schmidt, and Peer Bork. n.d. "GUNC: Detection of Chimerism and Contamination in Prokaryotic Genomes." https://doi.org/10.1101/2020.12.16.422776.

Parks, Donovan H., Maria Chuvochina, Christian Rinke, Aaron J. Mussig, Pierre-Alain Chaumeil, and Philip Hugenholtz. 2022. "GTDB: An Ongoing Census of Bacterial and Archaeal Diversity through a Phylogenetically Consistent, Rank Normalized and Complete Genome-Based Taxonomy." *Nucleic Acids Research*. https://doi.org/10.1093/nar/gkab776.

Parks, Donovan H., Michael Imelfort, Connor T. Skennerton, Philip Hugenholtz, and Gene W. Tyson. 2015. "CheckM: Assessing the Quality of Microbial Genomes Recovered from Isolates, Single Cells, and Metagenomes." *Genome Research* 25 (7): 1043–55. https://doi.org/10.1101/gr.186072.114.

Price, Morgan N., Paramvir S. Dehal, and Adam P. Arkin. 2010. "FastTree 2 – Approximately Maximum-Likelihood Trees for Large Alignments." *PLoS ONE*. https://doi.org/10.1371/journal.pone.0009490.

Ray, Angelique E., Julian Zaugg, Nicole Benaud, Devan S. Chelliah, Sean Bay, Hon Lun Wong, Pok Man Leung, et al. 2022. "Atmospheric Chemosynthesis Is Phylogenetically and Geographically Widespread and Contributes Significantly to Carbon Fixation throughout Cold Deserts.” *The ISME Journal* 16 (11): 2547–60. https://doi.org/10.1038/s41396-022-01298-5.

Seemann, Torsten. 2014. “Prokka: Rapid Prokaryotic Genome Annotation.” *Bioinformatics* 30 (14): 2068–69. https://doi.org/10.1093/bioinformatics/btu153.

Søndergaard, Dan, Christian N. S. Pedersen, and Chris Greening. 2016. “HydDB: A Web Tool for Hydrogenase Classification and Analysis.” *Scientific Reports* 6 (September): 34212. https://doi.org/10.1038/srep34212.

Tatusov, R. L., M. Y. Galperin, D. A. Natale, and E. V. Koonin. 2000. “The COG Database: A Tool for Genome-Scale Analysis of Protein Functions and Evolution.” *Nucleic Acids Research* 28 (1): 33–36. https://doi.org/10.1093/nar/28.1.33.
